# Emergence of Dual-Risk *Klebsiella pneumoniae*: Global Hotspots of Convergence Between Resistance and Hypervirulence

**DOI:** 10.1101/2025.11.13.688379

**Authors:** Prasanna Kumar Selvam, Supraja Mohan, Karthick Vasudevan

**Author notes:** **Corresponding:** Dr Karthick Vasudevan, Institute of Bioinformatics, International Technology Park, Bangalore, 560066, India.

## Abstract

**Background:** *Klebsiella pneumoniae* is a major pathogen causing both healthcare- and community-associated infections. Traditionally classified as classical (cKp) or hypervirulent (hvKp), recent reports suggest increasing overlap between antimicrobial resistance (AMR) and virulence, leading to “dual-risk” clones that are both invasive and difficult to treat.

**Objectives:** To perform a large-scale genomic analysis of *K. pneumoniae* isolates and systematically characterize the convergence of AMR and hypervirulence, with a focus on global distribution, clonal structure, and accessory genome functions.

**Methods:** We analyzed 3,443 genomes curated from 25 international studies. Virulence scoring was standardized using a Kleborate-based framework, while AMR genes were identified with ResFinder. Multilocus sequence typing (MLST), plasmid replicon typing, and K/O antigen profiling were conducted. Convergence was defined as hvKp (virulence score ≥3) carrying high-priority AMR determinants. Pan-genome-wide association (pan-GWAS) was performed with Roary and Scoary, followed by functional enrichment using ShinyGO.

**Results:** Among hvKp isolates, 59.4% carried carbapenemases, ESBLs, or 16S rRNA methyltransferases, demonstrating widespread convergence. Convergent lineages were dominated by ST231 and ST23, with India and the Middle East identified as geographic hotspots. HvKp isolates carried a higher plasmid replicon burden and were enriched in KL1/KL2 and O1/O2 serotypes. Pan-GWAS revealed hvKp-associated accessory genes linked to envelope stress response, nucleotide metabolism, and membrane transport, highlighting adaptive traits beyond canonical virulence loci.

**Conclusion:** Our findings redefine the hvKp/cKp dichotomy as a continuum shaped by plasmids and selective pressures. The emergence of dual-risk clones has urgent implications for diagnostics, surveillance, and treatment strategies in regions with high AMR prevalence.

## Introduction

The opportunistic Gram-negative pathogen *Klebsiella pneumoniae* is a growing global concern due to its remarkable genomic plasticity and clinical impact (Gomez-Simmonds & Uhlemann, 2017). Historically linked to nosocomial infections, it causes both invasive and healthcare-associated diseases (Al-Abeadi et al., 2023). Two major pathotypes are recognized: classical *K. pneumoniae* (cKp), associated with multidrug resistance (MDR) in hospital settings, and hypervirulent *K. pneumoniae* (hvKp), which causes severe community-acquired infections in otherwise healthy individuals (Liu et al., 2020). First identified in East Asia in the 1980s, hvKp is characterized by virulence plasmids and chromosomal loci encoding siderophores such as aerobactin (*iuc*), salmochelin (*iro*), and yersiniabactin (*ybt*), as well as hypermucoidy regulators (*rmpA*, *rmpA2*) (Lan & others, 2021). These factors promote systemic dissemination, immune evasion, and iron acquisition. While canonical hvKp strains are often antimicrobial-susceptible (Kochan & others, 2023), recent evidence shows increasing convergence between hvKp and cKp, with the acquisition of both AMR and virulence determinants (Russo & others, 2024; (WHO), 2024). This emergence of “dual-risk” clones complicates treatment strategies and poses significant infection control challenges.

The World Health Organization has reported the global spread of hypervirulent *K. pneumoniae* in 16 of 43 surveyed nations (Russo & Marr, 2019). Notably high prevalence has been documented in China (∼38%), France (∼20%), India (∼19%), Mexico (∼18%), Iran (∼15%), and Vietnam (∼14%) (Zhang & others, 2016) (Bhardwaj et al., 2020) (Yadav et al., 2023) (Nguyen & others, 2024). In India, the dual-risk phenotype is especially alarming, with up to 20% carbapenem resistance, >70% third-generation cephalosporin resistance, and ESBL rates approaching 80% (Sharma et al., 2023; Veeraraghavan & others, 2017). Since its discovery in Taiwan in 1986 (Marr & Russo, 2019) and the first carbapenem-resistant hvKp in India in 2016 (Shankar & others, 2016), hvKp has evolved from a regional entity to a global health priority. Recent outbreaks have identified carbapenemase genes (*bla*NDM, *bla*KPC) in hypervirulent ST23 and ST86 strains (Li & others, 2025), while MDR lineages such as ST231 and ST147 have acquired virulence plasmids carrying *pLVPK*-like elements (Du & others, 2021). These events challenge the binary hvKp/cKp classification and raise concerns over untreatable, highly invasive forms.

Understanding genomic distinctions between hvKp and cKp, and the mechanisms driving resistance–virulence convergence, is critical for public health planning (Choby et al., 2020). Most existing studies are limited in scale or lack standardized analytical frameworks. Here, we present one of the largest comparative genomic analyses of hvKp and cKp to date, integrating epidemiological metadata, sequence-based typing, gene content profiling, and trait association modeling. Our findings provide a resource for genomic surveillance, risk stratification, and the development of diagnostics, therapeutics, and control strategies.

## Materials and Methods

### Genome Collection

A total of 4,187 *K. pneumoniae* genomes were collected from publicly available databases, including NCBI, the Sequence Read Archive (SRA), and the European Nucleotide Archive (ENA), spanning 25 studies. The metadata accompanying these genomes such as country of origin and strain-level classification into hvKp or cKp were manually curated. Only those strains with clearly published phenotype classifications were retained, based on one or more of the following criteria: clinical presentation consistent with hvKp, positive string test indicating mucoviscosity, and/or PCR-based confirmation of key virulence genes (Figure 1). All accession IDs for the datasets used are listed in Supplementary Data.

**Figure 1.**
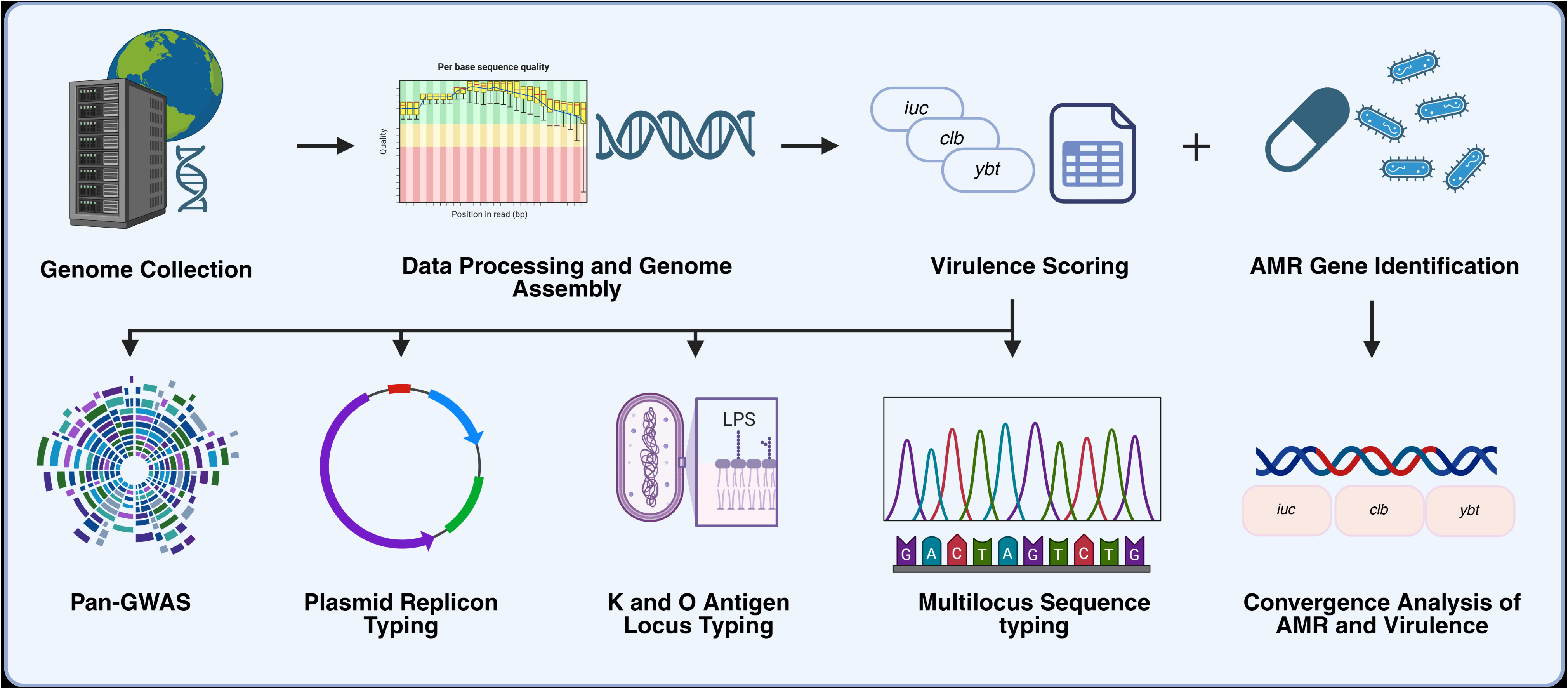
Overview of the genomic analysis pipeline for *K. pneumoniae*.

### Genome Assembly and Virulence Gene Scoring

Raw sequencing reads were obtained using fasterq-dump from the SRA Toolkit and quality-filtered with fastp v0.23.2 using default parameters. Unicycler v0.4.8 was used to construct the reads de novo. Genome assemblies were evaluated for quality using QUAST v5.2.0.

To address variability across 25 independent datasets, we applied a standardized virulence classification pipeline adapted from the Kleborate scoring system(Lam et al., 2021a). Virulence loci were detected using BLASTn (≥90% nucleotide identity, ≥80% coverage), with points assigned as follows: yersiniabactin (*ybt*), +1; colibactin (*clb*), +1; aerobactin (*iuc*), +2; salmochelin (*iro*, only if *iuc* present), +1; and *rmpA/rmpA2*, +1. Composite scores ranged from 0–5, with ≥3 classified as hvKp and <3 as cKp. Strains with discordant metadata and genomic classification were labeled as false hvKp/cKp and excluded from analysis.

### Multilocus Sequence Typing (MLST)

MLST analysis was performed to determine the clonal structure of the 3,443 *K. pneumoniae* genomes. Typing was conducted with Torsten Seemann’s mlst tool (https://github.com/tseemann/mlst)(v2.19.0), which employs the Pasteur 7-gene scheme to target the housekeeping genes *gapA, infB, mdh, pgi, phoE, rpoB,* and *tonB*. A local version of the *Klebsiella* MLST database, included with the mlst package, was used for allele identification and sequence type (ST) assignment, The distribution of STs between the hvKp and cKp groups allowed for lineage-level comparisons and the discovery of high-risk clones undergoing convergence.

### Antimicrobial Resistance Gene Detection

ABRicate (https://github.com/tseemann/abricatev) v1.0.1 was used to detect acquired AMR genes by searching assemblies against the ResFinder database. Hits were filtered using minimum limits of ≥90% nucleotide identity and ≥60% gene coverage. Resistance genes were classified into key functional classes, including β-lactams, carbapenemases, aminoglycosides, quinolones, and tetracyclines, based on their resistance patterns. The overall AMR gene load was computed for each genome. To evaluate resistance loading across hvKp and cKp pathotypes, we used the Wilcoxon rank-sum test, a non-parametric approach for determining differences in median gene counts between the two groups.

### Plasmid Replicon Typing

Plasmid replicon typing was carried out with ABRicate v1.0.1, which was compared to the PlasmidFinder database. The detection limits were established at ≥95% nucleotide identity and ≥60% minimum coverage. Replicon types found in each genome were recorded, and the frequency of important plasmid families was assessed across all isolates. A comparative study was then performed comparing the hvKp and cKp groups to assess variations in plasmid content and distribution.

### K and O Antigen Locus Typing

In silico typing of the capsular (K) and lipopolysaccharide (O) antigen loci was performed using Kaptive v2.0.3 to determine the serotype distribution across the *K. pneumoniae* dataset. Kaptive was run with default parameters. The prevalence of each K-locus (KL1 – KL170) and O-locus (O1v1 – O5) was computed separately for hvKp and cKp groups. The statistical significance of differences in locus frequencies between the groups was assessed using Fisher’s exact test, with *p* < 0.05 considered statistically significant.

### Convergence Analysis of AMR and Virulence

To explore genomic convergence between virulence and AMR, isolates were classified as convergent if they displayed a virulence score ≥3 (as indicated by the presence of *iuc*, *ybt*, *clb*, *iro*, *rmpA*/*rmpA2*) and simultaneously carried at least one high-priority AMR determinant. These determinants included carbapenemases, 16S rRNA methyltransferases, plasmid-mediated quinolone resistance genes, extended-spectrum β-lactamases (ESBLs), and colistin resistance markers, all identified using Abricate with the ResFinder database.

### Pan-genome Analysis and Accessory Gene Association

The assembled genomes then annotated using Prokka. The annotated GFF3 were used as input for Roary v3.13.0, with a BLASTp identity threshold of 95% and paralog splitting enabled. Gene clusters were classified as: Core genes: present in ≥99% of genomes, Soft-core: 95–99%, Shell: 15–95%, Cloud: <15%. Scoary v1.6.16 was used for pan-GWAS analysis to identify accessory genes significantly associated with the hvKp phenotype. The binary trait file used hvKp/cKp as classes based on the virulence score threshold. Genes with Bonferroni-corrected p-values <0.05 were considered significant.

### Functional Enrichment Analysis

We performed pathway enrichment on 200 hvKp-associated accessory genes identified using Scoary, prioritized by Youden’s Index (≥0.7), Positive and Negative Likelihood Ratios, and Benjamini–Hochberg FDR (<0.05). ShinyGO v0.76 (http://bioinformatics.sdstate.edu/go/) was used to assess enrichment against GO, KEGG, and other annotation databases, with FDR < 0.05 as the significance threshold. Enriched pathways were grouped into biologically relevant themes iron acquisition, envelope stress response, nucleotide biosynthesis, transport systems, DNA repair, and metabolic adaptation based on literature-supported hvKp virulence mechanisms and keyword-based pathway mapping.

## Results

### Sample Collection and Virulence Classification

A total of 4,187 *K pneumoniae* genomes were collected from publicly available sequencing projects spanning multiple countries, including China, India, Saudi Arabia, Germany, Egypt, Vietnam, and others. These genomes were curated from 25 studies and were initially annotated as either hvKp or cKp strains based on their associated metadata. Figure 2 summarizes the geographical distribution and strain classification of the *K. pneumoniae* genomes included in this study.

**Figure 2.**
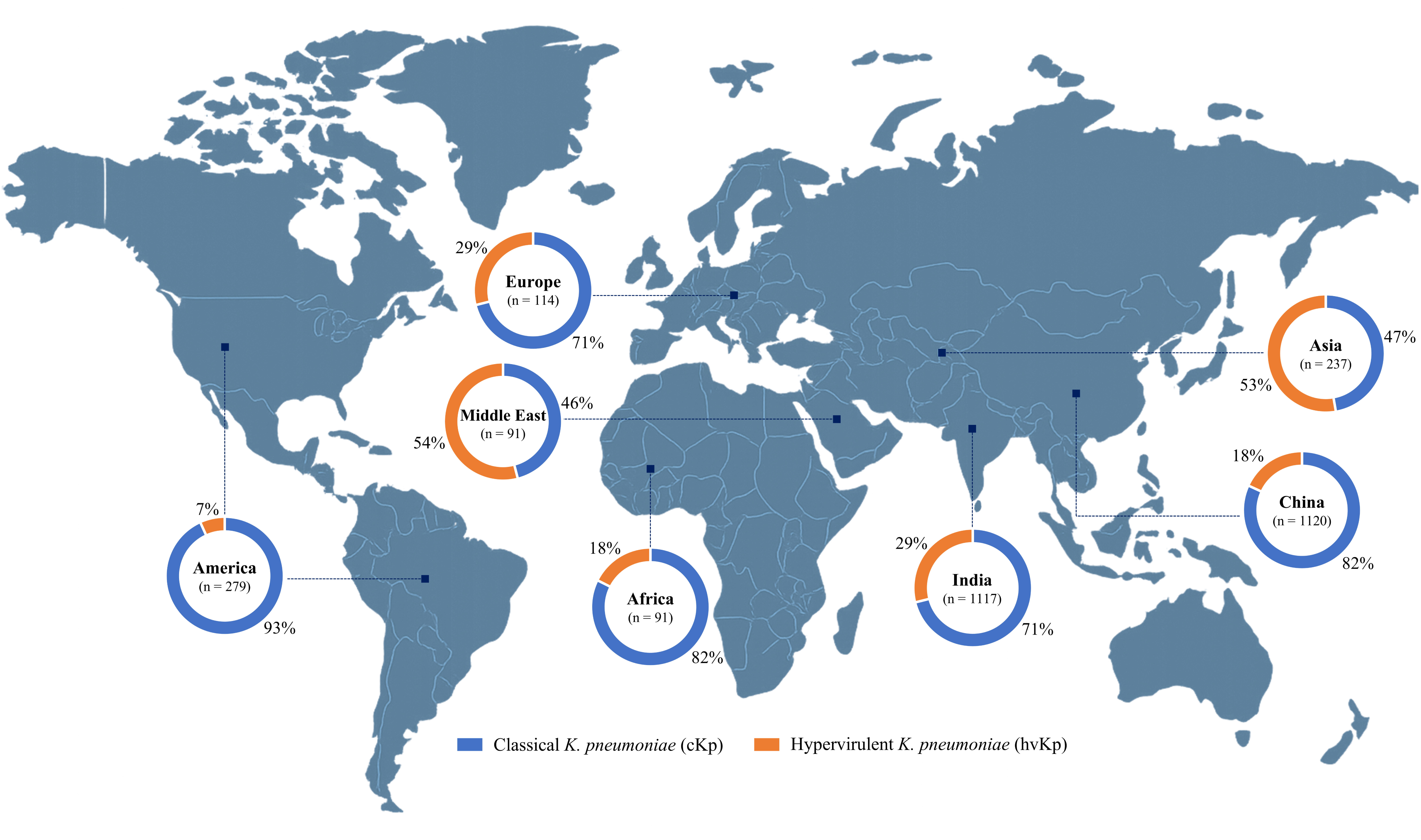
Global distribution and classification of 4,187 *K. pneumoniae* isolates.

To ensure uniform classification across diverse datasets, we implemented a virulence genotyping and scoring framework grounded in established criteria. Genomes were evaluated for the presence of key virulence loci: *yersiniabactin (ybt)* (+1), *colibactin (clb)* (+1), *aerobactin (iuc)* (+2), *salmochelin (iro)* (+1, only if *iuc* present), and *rmpA/rmpA2* (+1, only if *iuc* present). After reclassification with standardized virulence scoring, 980 hvKp and 2,463 cKp genomes were retained for downstream analyses (Supplementary Table 1).

### Multi-Locus Sequence Typing (MLST) Analysis

Out of the 3,443 *K. pneumoniae* isolates analyzed, 980 were classified as hvKp and 2,463 as cKp. Multi-locus sequence typing (MLST) revealed distinct clonal structures between the two groups (Fig. 3). HvKp isolates were dominated by ST231 and ST23, along with smaller contributions from ST65, ST86, and ST373. In contrast, cKp displayed broader diversity, with ST14, ST147, ST15, ST11, and ST395 being most common. Region-specific profiling showed that Indian hvKp isolates were largely associated with ST231, while Indian cKp were enriched in ST14 and ST147. Outside India, ST23 predominated among hvKp, whereas ST15 and ST11 were more frequent in cKp. These findings highlight clear lineage-level differences between hvKp and cKp as well as geographic variation in clonal distribution (Supplementary Table 1).

**Figure 3.**
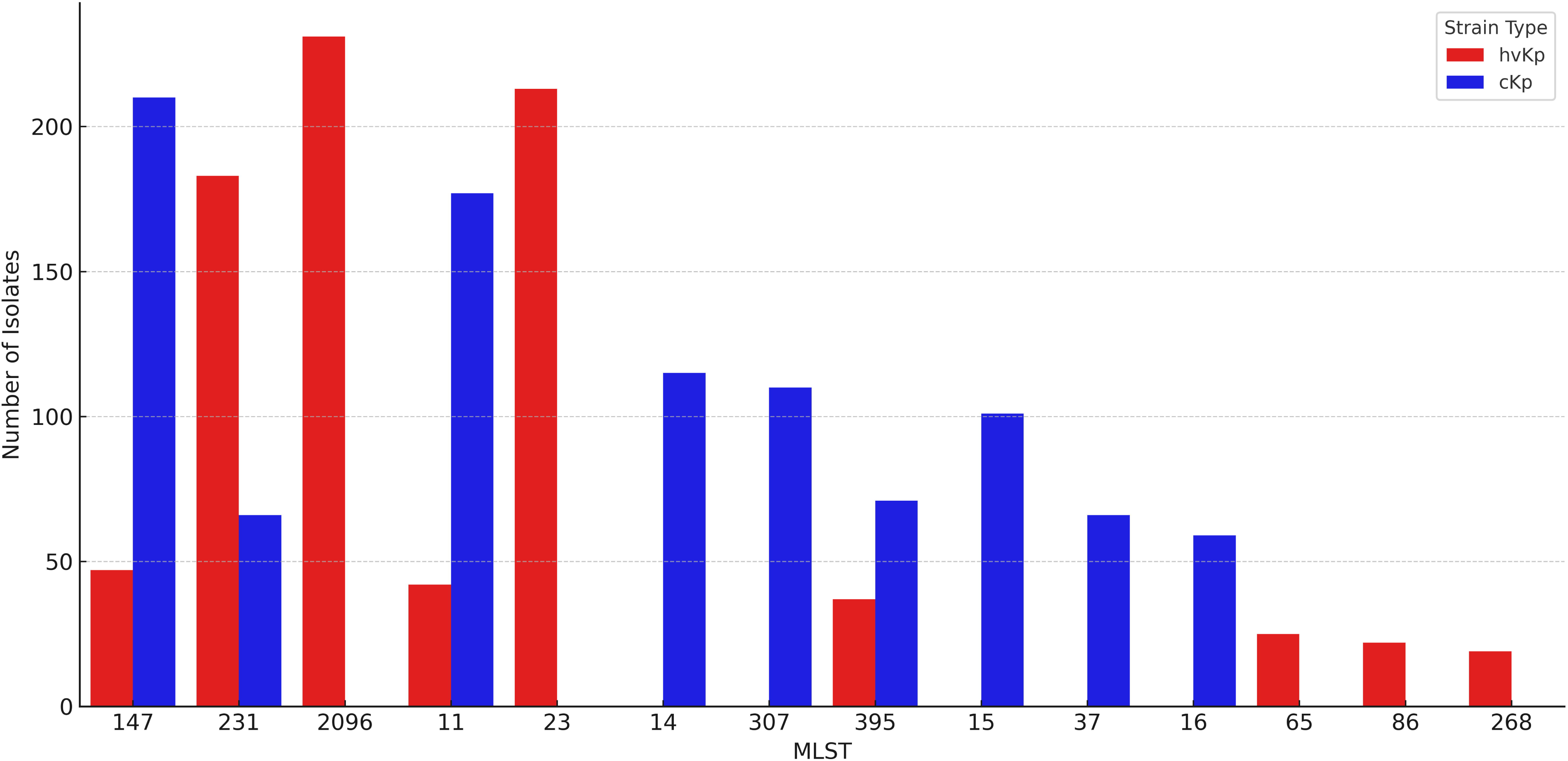
Distribution of major multi-locus sequence types (MLSTs) among *hvKp* and *cKp* isolates. Bar plot showing the number of *K. pneumoniae* isolates belonging to the most prevalent sequence types (STs), stratified by strain type: hypervirulent (*hvKp*, red) and classical (*cKp*, blue). While *hvKp* was dominated by ST231 and ST23, *cKp* exhibited a broader ST diversity, with ST14 and ST147 among the most common.

### Antimicrobial Resistance Gene Profiling

To characterize the AMR landscape of 3,443 *K. pneumoniae* isolates, ResFinder was used to identify acquired resistance genes (≥90% identity, ≥60% coverage). Nearly all isolates (98.4%) carried at least one resistance determinant, with widespread evidence of multidrug resistance. β-lactamase genes were most common, led by *bla*CTX-M (64.3%), followed by blaSHV and *bla*TEM. Carbapenemase genes, including *bla*NDM-1 (12.7%), *bla*OXA-232 (9.6%), and *bla*KPC-2 (6.1%), were detected predominantly in cKp, though occasional acquisitions in hvKp signaled convergence of resistance and virulence. Aminoglycoside resistance genes (*aac(3)-IIa, aadA1, aph(3’)-Ia*) occurred in 47.5% of isolates, while plasmid-mediated fluoroquinolone genes (*qnrB, qnrS1*), tetracycline genes (*tetA, tetM*), and sulfonamide genes (*sul1, sul2*) were also frequent.

Overall, cKp carried a significantly higher AMR burden than hvKp (median 11 vs. 4 genes; p < 0.0001). Indian isolates showed markedly greater AMR load compared to global counterparts (median 15 vs. 7 genes; p = 4.22 ×10¹), with cKp frequently harboring *bla*NDM-1 and *bla*OXA-232, and hvKp occasionally acquiring carbapenemases alongside virulence loci. These findings underscore distinct resistance profiles between hvKp and cKp and highlight India as a hotspot for resistance–virulence convergence. The prevalence of major AMR genes and country-wise gene burden is shown in Fig. 4A–B (Supplementary Table 2).

**Figure 4A.**
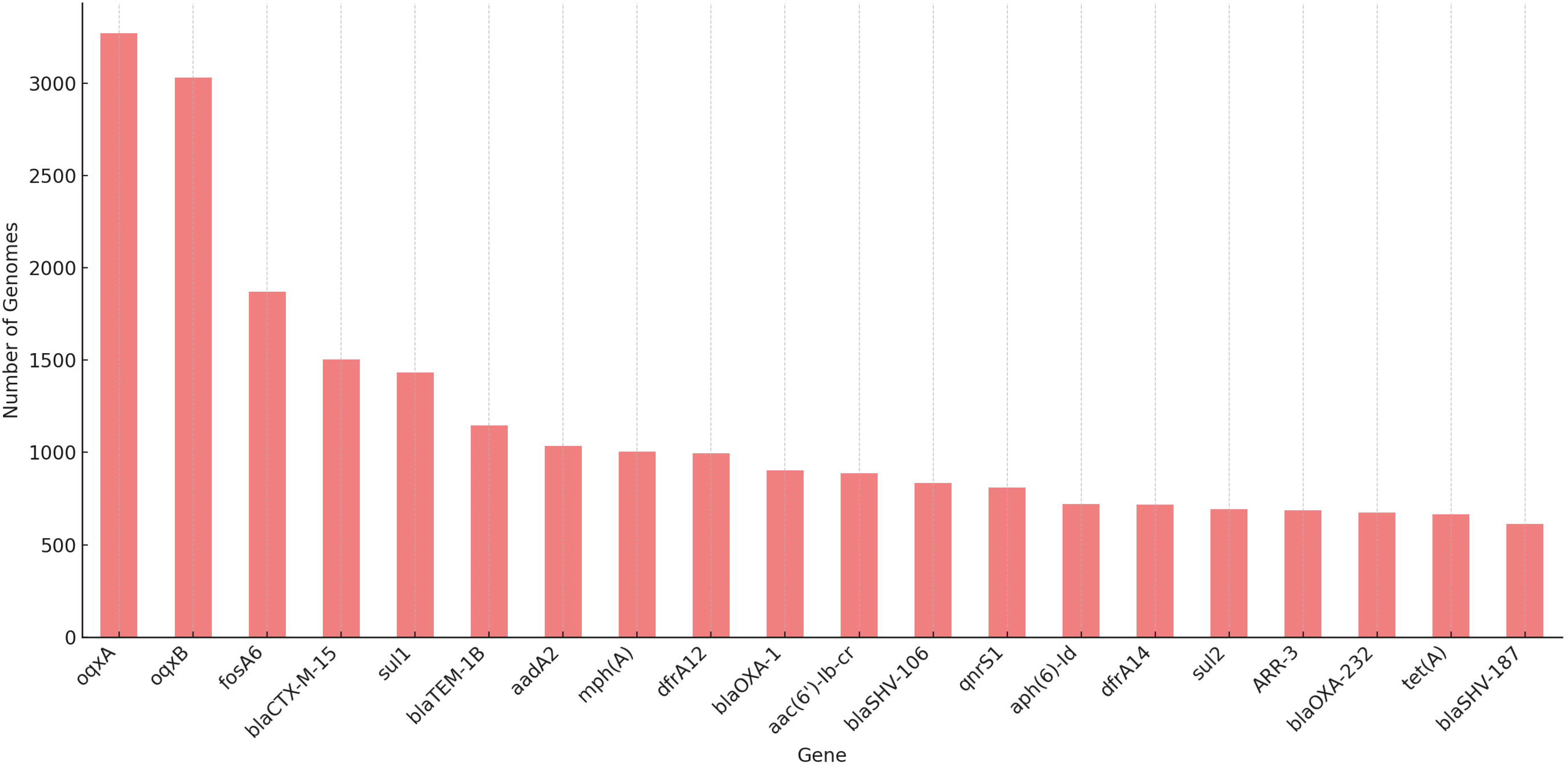
Prevalence of major antimicrobial resistance genes in *K. pneumoniae* genomes. Bar chart showing the number of genomes carrying each of the top 25 most frequently detected AMR genes. Genes conferring resistance to β-lactams (*blaCTX-M-15*, *blaSHV*, *blaTEM-1B*), aminoglycosides (*aadA2*, *aph(6)-Id*), fluoroquinolones (*qnrS1*), and sulfonamides (*sul1*, *sul2*) were dominant across the genomes.

**Figure 4B.**
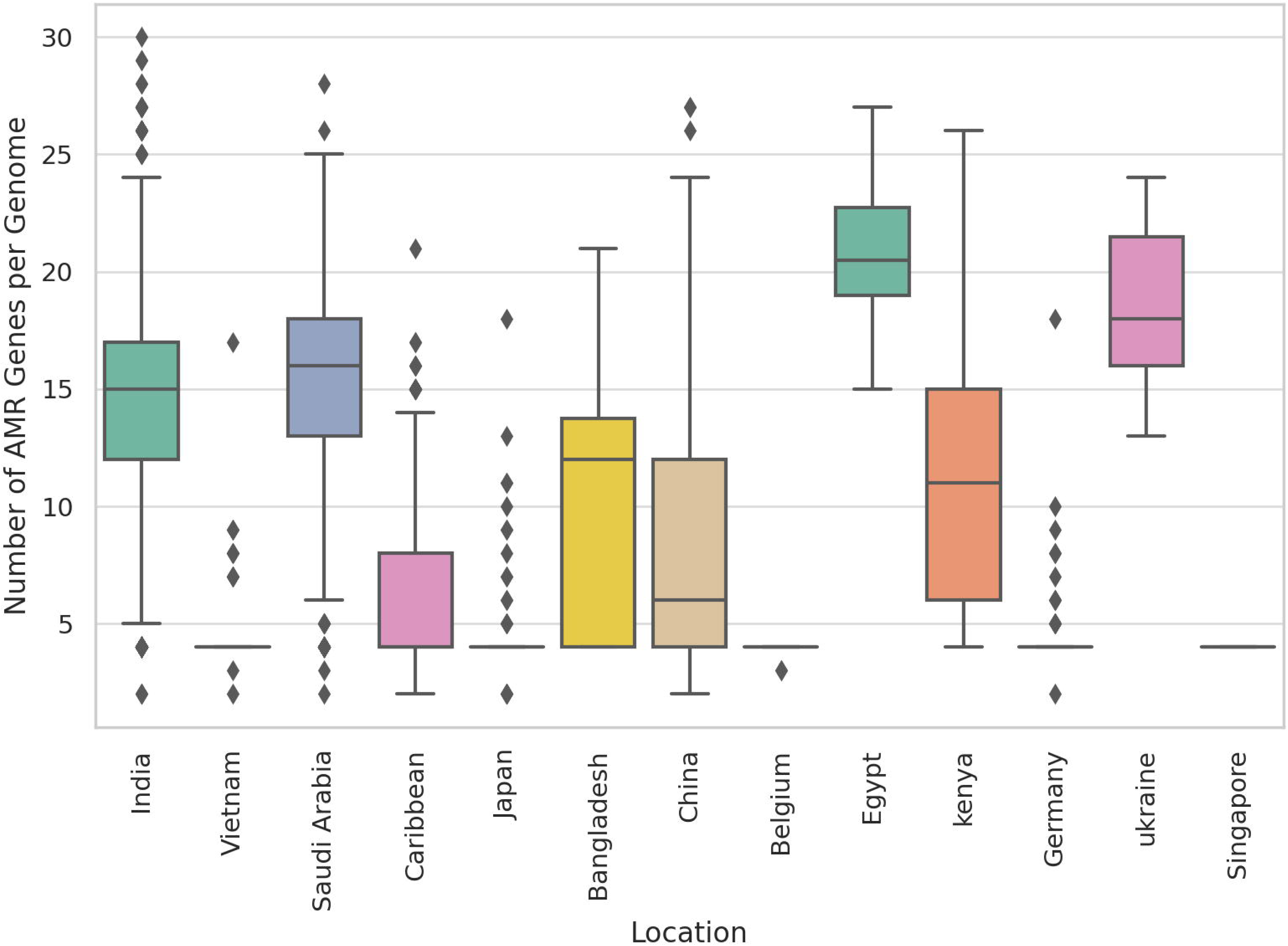
Geographic distribution of AMR gene burden in *K. pneumoniae*. Boxplot depicting the number of acquired AMR genes per genome across different countries. Indian isolates exhibited the highest median AMR gene burden, followed by Germany, Egypt, and Saudi Arabia. The data reflect regional disparities in resistance gene prevalence and potential hotspots of multidrug resistance.

### Plasmid Replicon Profiles Differentiate hvKp and cKp Isolates

Plasmid replicon profiling of 3,443 *K. pneumoniae* isolates using PlasmidFinder showed that 93% carried at least one plasmid replicon. The number of replicons per genome varied widely (median 4; range 1–17). HvKp isolates harbored a significantly broader plasmid repertoire than cKp (median 5 vs. 3 replicons per genome; p = 3.27 × 10², Wilcoxon test), consistent with their increased capacity for horizontal gene acquisition.

Geographic comparisons revealed a similar trend, with Indian isolates carrying a higher plasmid burden (median 5 vs. 3 replicons in global isolates; p = 2.04 × 10¹). This enrichment highlights India as a hotspot for diverse plasmid types that facilitate the co-occurrence of resistance and virulence determinants (Supplementary Table 3).

### K and O Antigen Profiling of *K. pneumoniae* Isolates

In silico serotyping of 3,443 genomes identified 72 K loci and 12 O loci. HvKp isolates were dominated by KL1 and KL2, alongside O1/O2 serotypes, whereas cKp displayed broader diversity, including KL102, KL107, KL24, and O3b/O5. Enrichment of O1/O2 variants in hvKp was statistically significant (p < 0.001, Fisher’s exact test). Among Indian isolates, hvKp was similarly restricted to KL1/KL2, while cKp showed greater antigenic diversity. Overall, hvKp exhibited limited serotype distribution, reinforcing its clonal dominance, whereas cKp remained more antigenically diverse (Supplementary Table 1).

### Convergence of Antimicrobial Resistance and Hypervirulence

Of the 3,443 isolates, 64.3% carried at least one high-priority AMR gene. Convergent hvKp accounted for 17% of all isolates and 59% of hvKp strains, underscoring the growing overlap between resistance and virulence. These lineages were concentrated in India (48%) and Saudi Arabia (44%), with smaller clusters in China and Vietnam. In contrast, most AMR-positive cKp isolates had low virulence scores, indicating that true convergence is largely confined to hvKp. Figure 5 illustrates both the higher AMR burden in hvKp compared to cKp (5A) and the geographic clustering of convergent hvKp (5B).

**Figure 5A.**
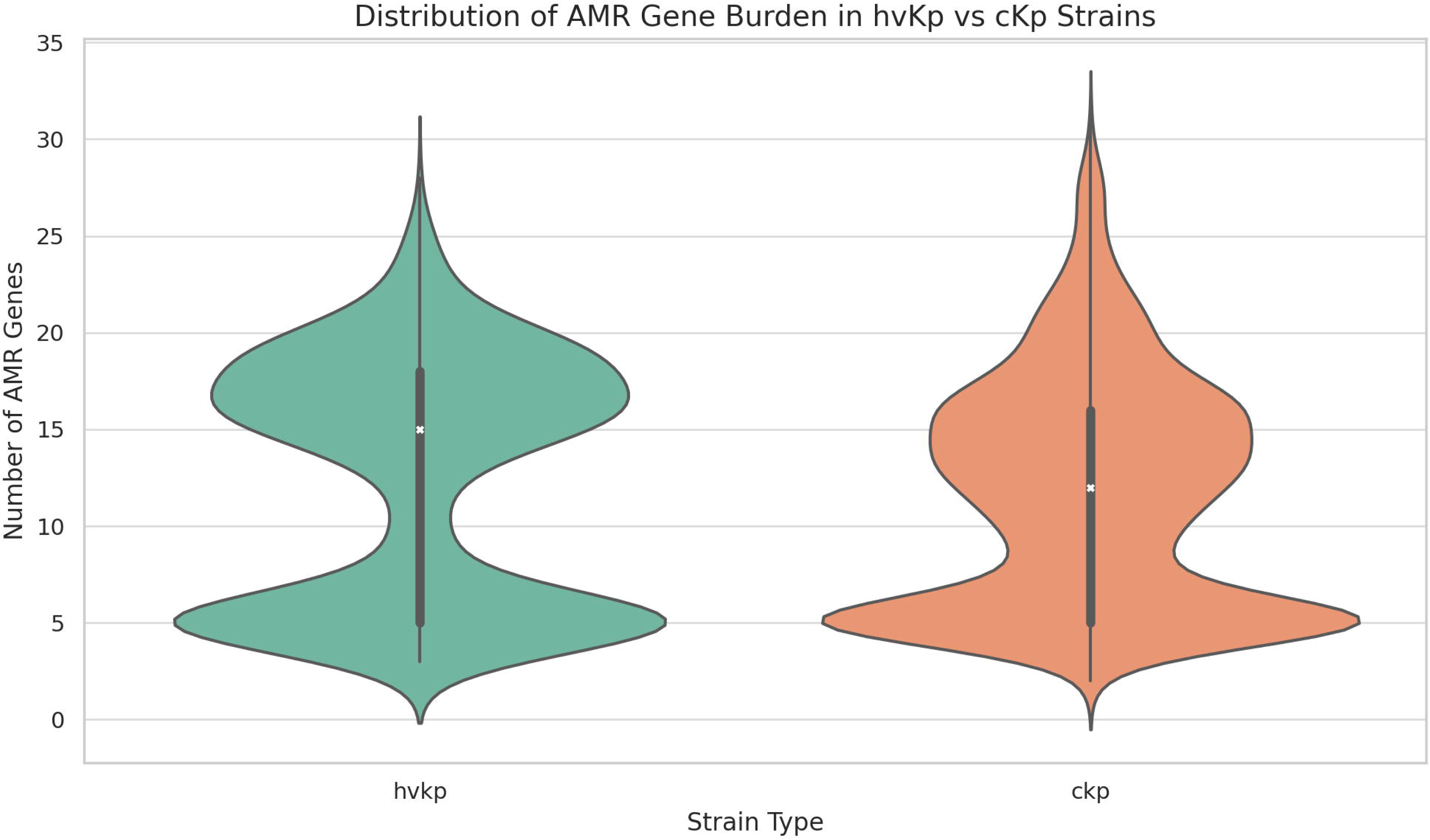
Distribution of AMR gene burden in hvKp vs cKp strains. Violin plot comparing the number of acquired AMR genes between hypervirulent (*hvKp*) and classical (*cKp*) *K. pneumoniae* isolates. While both lineages carry resistance genes, hvKp strains tend to exhibit a bimodal distribution with a lower median AMR burden compared to cKp.

**Figure 5B.**
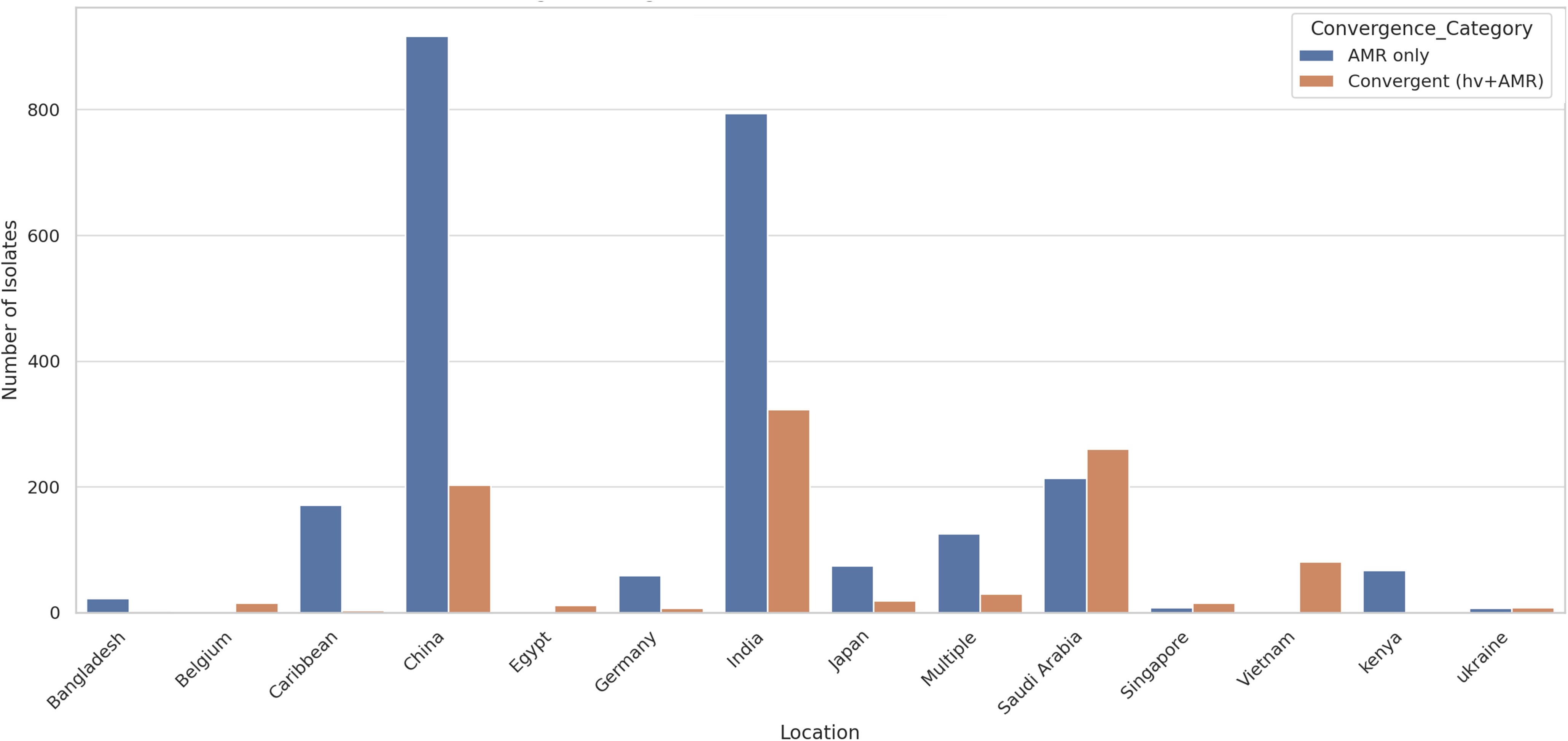
Geographic distribution of convergent hvKp isolates. Bar chart showing the number of *K. pneumoniae* isolates with convergence of AMR and virulence traits (orange) compared to those carrying AMR genes only (blue), across different geographic locations. India and Saudi Arabia represent the highest contributors of convergent hvKp, suggesting regional hotspots of resistance–virulence overlap.

### Genome Annotation and Pan-genome Construction

All 3,443 *K. pneumoniae* genomes were uniformly annotated using Prokka v1.14.6, generating standardized GFF3 files for pan-genome construction. Pan-genome analysis was performed using Roary v3.13.0, resulting in a total of 135,612 orthologous gene clusters. Of these, 3,241 genes were classified as part of the core genome (present in ≥99% of isolates), while the remaining 132,371 genes comprised the accessory genome, including soft-core (95–99%), shell (15–95%), and cloud genes (<15%). This large accessory genome highlights the high genomic plasticity of both hvKp and cKp lineages.

### Pan-GWAS to Identify Hypervirulence-Associated Genes

Pan-genome-wide association analysis using Scoary identified a distinct set of accessory genes linked to the hvKp phenotype (Bonferroni-corrected p < 0.05). As expected, canonical virulence loci including aerobactin (*iuc*), salmochelin (*iro*), and the regulators *rmpA/rmpA2* were strongly enriched, validating the analytical approach. Beyond these, hvKp genomes showed significant enrichment of mobile genetic elements such as insertion sequences (*IS903, ISYps3*) and transposon-related genes (e.g., *Tn3*), many located in proximity to virulence loci. These findings highlight the role of horizontal gene transfer in shaping the hvKp accessory genome and suggest that mobile elements contribute to the mobilization and dissemination of pathogenicity islands (Supplementary Table 4).

### Functional enrichment analysis

Functional enrichment of the top 200 hvKp-associated accessory genes (prioritized by Youden’s Index via Scoary) was assessed in ShinyGO against the *K. pneumoniae* genome background (FDR < 0.05). The top ten pathways, spanning envelope stress response, metabolic adaptation, nucleotide biosynthesis, and membrane transport, included the rpoE–rseB–mucB regulon (FDR = 0.0026, fold enrichment = 31.76), protein folding/redox regulation genes (LSPA, FKPB, YHAI; FDR = 0.0053, fold = 47.65), lipid metabolism/isoprene biosynthesis (ISPH; FDR = 0.0079, fold = 18.68), nucleotide biosynthesis/kinase activity (NRDB, GUAD_1, RIHC, CARB; FDR = 0.0079, fold = 5.10), and nucleoside H symporters (HCAT, NUPG; FDR = 0.0435, fold = 52.94). All were supported by Bonferroni-adjusted Scoary results and mapped to hvKp-associated clusters across diverse lineages (Figure 6; Supplementary Table 5).

**Figure 6.**
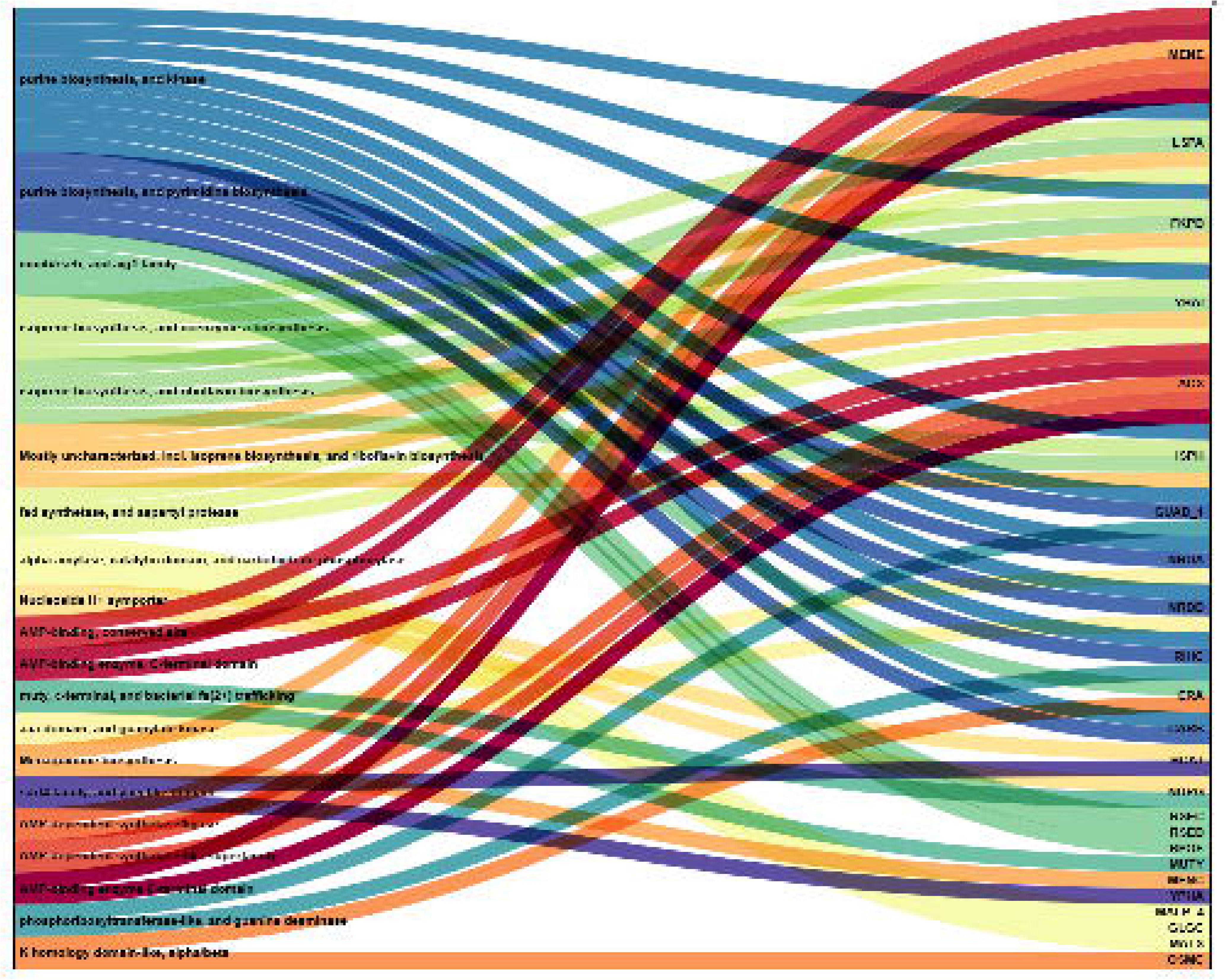
Alluvial plot of the top 20 enriched pathways among hvKp-associated accessory genes. Alluvial plot showing the mapping between the top 20 significantly enriched functional pathways (left) and their corresponding gene annotations (right) among the top 200 accessory genes associated with hypervirulent *K. pneumoniae* (hvKp). Pathways were identified using ShinyGO enrichment analysis (FDR < 0.05) and represent diverse functional modules, including envelope stress response, redox regulation, nucleotide biosynthesis, isoprene/lipid metabolism, and membrane transport. The width of each flow corresponds to the number of genes contributing to each pathway. Genes shared across multiple pathways are represented by flows branching from different categories.

## Discussion

This study represents one of the largest comparative genomic analyses of hvKp and cKp to date, integrating population structure, virulence architecture, AMR, and convergence patterns into a unified framework. By standardizing virulence scoring through an in-house Kleborate-inspired framework (Lam et al., 2021b), we resolved inconsistencies in pathotype classification from 25 global datasets, enabling accurate comparisons of regional and lineage-specific trends in resistance and antigenic diversity. To our knowledge, this is among the most comprehensive assessments of molecular overlap between hvKp and cKp, underscoring the emerging threat of dual-risk convergent clones.

Traditionally, hvKp has been linked to community-acquired invasive infections with low AMR, while cKp predominates in nosocomial settings with high resistance burdens. Our findings challenge this binary paradigm: 59.4% of hvKp isolates carried at least one high-priority AMR determinant, including carbapenemases, ESBLs, and 16S rRNA methyltransferases. This rapid convergence with multidrug-resistant (MDR) backgrounds aligns with reports from East and South Asia of hvKp strains harboring *bla*NDM-1, *bla*OXA-232, and *bla*KPC-2. Convergence was most frequent in high-risk sequence types ST231 and ST23, commonly associated with pLVPK-like virulence plasmids and IncFIB(K) replicons that facilitate co-localization of resistance and virulence genes (Choby et al., 2020; Tang et al., 2020).

Geographic stratification showed convergence concentrated in India and Saudi Arabia, reflecting strong antibiotic selection pressures and limited infection control. In India, the dominance of *bla*NDM-1, *bla*OXA-232, and IncFIB(K)/IncHI1B plasmid backbones mirrors national surveillance data, supporting India’s role as a global epicenter of AMR–virulence convergence (Shankar & others, 2022). The enrichment of ST231 among Indian hvKp, frequently carrying aerobactin (*iuc*), *rmpA2*, and multiple plasmid replicons, highlights its rise as a globally disseminating convergent lineage implicated in outbreaks across Southeast Asia and the Middle East (Shaidullina & others, 2023).

Plasmid profiling indicated that hvKp harbors significantly more replicons than *c*Kp (median 5 vs 3), suggesting a greater capacity for horizontal gene acquisition. HvKp harbored more plasmid replicons than cKp, highlighting their enhanced capacity for horizontal gene acquisition. Indian and Middle Eastern isolates were particularly enriched in IncHI1B and IncFIB plasmids, key vehicles for convergence (Bethke et al., 2023; Hammad et al., 2025).

Serotyping and virulence genotyping confirmed the dominance of KL1/KL2 and O1/O2 serotypes in hvKp, whereas cKp exhibited broader antigenic diversity. Notably, some high-AMR cKp isolates (e.g., ST15, ST147, ST11) carried partial virulence loci such as *ybt* or *rmpA2*, representing potential “cryptic” intermediates along the convergence pathway with the capacity to acquire additional virulence traits under selective pressure (Martin & others, 2023).

Pan-genome analysis highlighted the remarkable genomic plasticity of *K. pneumoniae*, with extensive accessory gene diversity distinguishing hvKp from cKp. HvKp genomes were enriched in canonical virulence loci such as *iuc*, *iro*, and *rmpA/rmpA2*, along with mobile genetic elements including insertion sequences and transposons. Many of these elements were located near virulence clusters, supporting their role in mobilizing and disseminating pathogenicity islands such as ICEKp10 and the pLVPK plasmid. These findings underscore the importance of horizontal gene transfer in shaping hvKp evolution and driving convergence with resistant lineages (Han & others, 2025).

Functional enrichment analysis of hvKp-associated accessory genes revealed significant representation of pathways critical for bacterial stress tolerance, metabolism, and host adaptation. The envelope stress response was the most enriched category, including *rpoE*, *rseB*, and *rseC*, key regulators of the σEregulon that maintains outer membrane integrity under osmotic and oxidative stress (Hews et al., 2019). Activation of this regulon enhances periplasmic folding and chaperone production, aiding survival against neutrophil-derived ROS and complement-mediated lysis (Fan et al., 2022). The presence of *mucB* homologs supports a role in anti-sigma factor regulation, while *fkpB* suggests chaperone-mediated stability of virulence secretion systems under host stress (Flores-Kim & Darwin, 2014; Tong & Jiang, 2015).

Isoprenoid biosynthesis and coenzyme metabolism, notably the *isph* gene of the non-mevalonate (MEP) pathway, were also enriched. This pathway is essential for undecaprenyl phosphate production, a lipid carrier required for peptidoglycan and LPS assembly (Manat & others, 2014). Its absence in humans makes it a potential hvKp-specific drug target. Additionally, purine biosynthesis and kinase activity modules (*guaD_1*, *ndb*, *rihC*, *carB*) were enriched, supporting nucleotide homeostasis and virulence gene expression in nutrient-limited tissues such as the liver and spleen (Acierno & others, 2025; Samant & others, 2008).

Membrane transporters such as *hcaT* and *nupG* (nucleoside:H<+symporters) were enriched, likely conferring an advantage in scavenging host-derived nucleosides and maintaining nucleotide pools in low-availability environments such as the bloodstream (Wang & others, 2021). These functional traits, coupled with canonical virulence loci, provide hvKp with metabolic flexibility and enhanced stress resilience, supporting persistence, dissemination, and immune evasion. Their frequent linkage to mobile elements suggests modular acquisition, making them attractive antivirulence targets (Martínez et al., 2019).

Overall, resistance hypervirulence convergence in *K. pneumoniae* is neither rare nor sporadic but a rapidly expanding phenomenon driven by plasmid exchange, mobile elements, and regional selection pressures. Convergent hvKp accounted for 17% of all isolates and 59% of hvKp, surpassing prior estimates. While *c*Kp remains dominant, the growing exchange of resistance and virulence traits signals converging evolutionary trajectories toward highly invasive, treatment-resistant strains a genomic “perfect storm”. Clinically, these genomic features are concerning because dual-risk clones complicate empiric therapy: hypervirulence combined with carbapenem resistance leaves few effective treatment options. Conventional AST, capsule typing, or clinical presentation alone may fail to identify such high-risk hybrids, underscoring the need for integrated genomic diagnostics that capture both virulence and resistance determinants.

Our findings emphasize the need for genomic surveillance that integrates virulence genotyping with AMR profiling, as conventional AST, capsule typing, or clinical presentation alone may overlook emerging high-risk hybrids. This genomic blueprint highlights key convergence markers suitable for incorporation into rapid diagnostics to detect virulence and resistance simultaneously. Such approaches, combined with targeted infection control, may help contain the spread of dual-risk clones. This study is limited by reliance on publicly available genomes, which may underrepresent certain geographic regions, and by short-read sequencing, which restricts resolution of plasmid architectures critical for resistance–virulence convergence. Future work using long-read sequencing and prospective surveillance will be essential to validate and expand these findings.

### Conclusion

Our global comparative analysis of 3,443 *K. pneumoniae* genomes reveals extensive genomic convergence between hypervirulence and antimicrobial resistance, with 59.4% of hvKp strains carrying high-priority AMR genes. This convergence, largely driven by plasmid exchange and mobile genetic elements, is particularly pronounced in ST231 and ST23 lineages from India and the Middle East. Functional enrichment highlights accessory genes involved in envelope stress response, metabolism, and host adaptation as key contributors to the hvKp phenotype. These findings challenge the conventional hvKp/cKp dichotomy and highlight the rise of dual-risk clones combining high virulence and multidrug resistance. Our study emphasizes the urgent need for integrated genomic surveillance to monitor, track, and mitigate the global spread of these increasingly untreatable pathogens.

## Authorship contribution

**P.K.S:** Writing – review & editing, Writing – original draft, Visualization, Validation, Methodology, Formal analysis, Data curation, Conceptualization. **S.M:** Writing – review & editing, Formal analysis. **K.V:** Writing – review & editing, Project administration, Supervision.

## Data Availability

Genome assemblies and metadata used in this study are available from NCBI/ENA/SRA under accession numbers listed in Supplementary Table 1.

## Ethical Approval

This study was based solely on publicly available genomic data. No ethical approval was required

## Acknowledgments

The authors express deep gratitude to the management of Manipal Academy of Higher Education (MAHE) and Institute of Bioinformatics (IOB), for providing all the support, necessary facilities, assistance, and constant encouragement to carry out this work.

